# Site Effects in Multisite Fetal Brain MRI: Morphological Insights into Early Brain Development

**DOI:** 10.1101/2023.12.16.572004

**Authors:** Xinyi Xu, Cong Sun, Hong Yu, Guohui Yan, Qingqing Zhu, Xianglei Kong, Yibin Pan, Haoan Xu, Tianshu Zheng, Chi Zhou, Yutian Wang, Jiaxin Xiao, Ruike Chen, Mingyang Li, Songying Zhang, Hongjie Hu, Yu Zou, Jingshi Wang, Guangbin Wang, Dan Wu

## Abstract

**Background:** Multisite MRI studies have become prevalent given their advantage in revealing reliable biological or clinical findings. Adult and adolescent multisite studies have demonstrated inevitable site-related non-biological effects that introduce bias. However, the site effect on fetal brain MRI remains unknown.

**Purpose:** To identify crucial acquisition factors affecting fetal brain structural measurements and developmental patterns, while assessing the effectiveness of existing harmonization methods in mitigating site effects.

**Materials and Methods:** Between May 2017 and March 2022, T2-weighted fast spin-echo sequences in-utero MRI were performed in healthy fetuses from prospectively recruited pregnant volunteers on four different scanners at four sites. A generalized additive model (GAM) was used to quantitatively assess site effects, including field strength (FS), manufacture (M), in-plane resolution (R), and slice thickness (ST), on subcortical volume and cortical morphological measurements, including cortical thickness, curvature, and sulcal depth. Growth models were selected to elucidate developmental trajectories of these morphological measurements. Welch’s test was performed to evaluate the influence of site effects on developmental trajectories. ComBat-GAM harmonization method was applied to mitigate site-related biases.

**Results:** The final analytic sample consisted of 340 MRI scans from 218 fetuses (mean gestational age, 30.1 weeks ± 4.4 [range, 21.7–40 weeks]). GAM results showed that low FS and low spatial resolution led to overestimations in selected brain regions of subcortical volumes and cortical morphological measurements, and cortical measurements were more susceptible to site effects than subcortical volumes. Only the peak cortical thickness in developmental trajectories was significantly influenced by the effects of FS and R. Notably, ComBat-GAM harmonization effectively removed site effects while preserving developmental patterns.

**Conclusion:** Our findings pinpointed the key acquisition factors in in-utero fetal brain MRI and underscored the necessity of data harmonization when pooling multisite data for fetal brain morphology investigations.

## 1 Introduction

In-utero fetal MRI has emerged as a crucial diagnostic tool for evaluating normal and pathological brain development[1]. High-resolution imaging of the fetal brains in utero can quantitatively characterize the spatiotemporal developmental trajectory of the fetal brain and thus provide the normative template for identifying high-risk fetuses before birth [2]. However, obtaining fetal brain MRI is often challenging due to the challenges in image acquisition related to fetal and maternal motions, patient recruitment, and the additional complexity of volumetric reconstruction required before quantification[3], resulting in smaller sample sizes and inconsistent findings in fetal brain MRI studies[4–9].

It is a common practice to pool MRI data acquired from different centers in order to increase the statistical power [10]. Large scale consortia, such as Human Connectome Project (HCP, https://www.humanconnectome.org/)[11], Enhancing Neuro Imaging Genetics through Meta-Analysis (ENIGMA)[12], UK Biobank (https://www.ukbiobank.ac.uk/)[13], have been established to aggregate large-cohort multicenter neuroimaging data to improve the reliability and reproducibility of neuroscience research. A major concern when analyzing results from these large consortium datasets is the potential bias introduced by non-biological site-related effects[14]. Previous studies have demonstrated that the site effects, including field strength, manufacture, software upgrade, as well as acquisition protocols, introduced bias and variance in measurements of brain volume[15–17], cortical thickness[18–23], functional and diffusion MRI[24–30] in adults, adolescents, children, infants and neonates. Nevertheless, the impact of these site effects on fetal brain MRI remains unknown.

To address the issue of site effects, researchers have proposed various harmonization methods to remove site-related variabilities while preserving the underlying biological properties in the data. For example, ComBat, which employs empirical Bayes framework to improve the estimation of the site-specific scaling parameters[31], has been commonly used to remove site-related effects while preserving biological variations. ComBat has demonstrated its effectiveness in harmonizing structural MRI measurements, including subcortical volume and surface area[14], cortical thickness[18], as well as diffusion MRI [32] and functional MRI[33] related measurements. Subsequently, Pomponio et al. introduced the ComBat-GAM harmonization method, which incorporated a generalized additive model (GAM) to capture nonlinear age effects across the lifespan[34].

In this study, we aim to investigate the influence of key acquisition factors on subcortical brain volume and cortical morphology descriptors in normal fetal brains, using 340 in-utero T2-weighted MRI scans of healthy fetal brains obtained from four different sites. Additionally, we explored whether scanner-related effects could be effectively mitigated using state-of-the-art harmonization method for robust and cross-center characterization of normal fetal brain development. We also generated spatiotemporal atlases of the fetal brain at 1.5T and 3T (acquired with thin/thick slices) separately to serve normative templates for different purposes.

## 2 Material and Methods

### 2.1 Data acquisition and subjects

The T2-weighted fetal brain MRI data were acquired from healthy fetal brains across four different hospitals. All Ethical approval was obtained from the Institutional Review Board at each participating hospital, and all participants provided written informed consent. All fetal brain MRI scans were obtained using T2-weighted images in orthogonal planes (axial, coronal, and sagittal), which is a common approach to correct fetal (and maternal) inter-slice motion and generate 3D isotropic volumes for quantitative analysis [35]. No maternal sedation was used in any of the scans. The detailed acquisition protocol was described in the following and the major site-specific acquisition factors are summarized in Table 1. Fetuses with multiple gestational pregnancies, intrauterine growth restriction, or other identified brain abnormalities were excluded from this study. Fetuses with ventriculomegaly, visible motion artifacts on MRI, or failed subsequent data processing were also excluded. In the end, 340 scans from 218 healthy fetal brains with gestational age (GA) from 21.7-40 weeks (30.1±4.4 w) were included in the final analysis. A participant flowchart is presented in Figure S1.

**Table 1.** Key acquisition parameters of fetal brain T2w MRI images in the five datasets.

| Site Name | Location | Field Strength (T) | Manufacturer | Age | In-plane Resolution (mm<sup>2</sup>) | Slice thickness (mm) | No. Participants | Dataset |
| --- | --- | --- | --- | --- | --- | --- | --- | --- |
| *Shandong Provincial Hospital* | Jinan | 3 | Siemens | 21.7 - 40w | 1.09×1.09 | $\frac{2}{4}$ | 122 | $\frac{1}{2}$ |
| *Dalian Municipal Women and Children's Medical Center* | Dalian | 1.5 | GE | 23.8 - 34w | 1.64 or 1.56 | 4 | 48 | 3 |
| *Women's Hospital of Zhejiang University Medical School* | Hangzhou | 1.5 | GE | 23 - 37w | 0.74 or 0.7031 | 2-5 | 13 | 4 |
| *Sir Run Run Shaw Hospital* | Hangzhou | 1.5 | Siemens | 22 - 38w | 1.02×1.02 | 4 | 35 | 5 |

#### Dataset 1&2 – Shandong Provincial Hospital (SPH)

We acquired MRI from 181 pregnant women at Shandong Provincial Hospital (SPH) between October 2019 and September 2021, with GA ranging from 20 to 40 weeks. After exclusion, 122 healthy singleton fetuses at GAs between 21.7 and 40 weeks were included in the following analysis. All fetuses were scanned on a 3.0 T Siemens scanner (MAGNETOM Skyra, Siemens Healthcare, Germany), using a Half-Fourier Acquisition Single-shot Turbo spin Echo (HASTE) imaging sequence in multiple axial, coronal, and sagittal orientations. The imaging parameters were as follows: repetition time (TR) = 800 ms, echo time (TE) = 113 ms/ 97 ms, no slice gap, in-plane resolution = 1.09×1.09 mm^2^. Notably, each fetus underwent two sets of MRI scans at 2 mm and 4 mm slice thickness, which were considered as two separate datasets in the subsequent analysis.

#### Dataset 3 – Dalian Municipal Women and Children’s Medical Center (DWC)

84 healthy volunteers at 23.6 to 37 weeks GA were recruited at Dalian Municipal Women and Children’s Medical Center (DWC) from May 2017 to December 2020. After quality check, 48 subjects at 23.8w to 34w GA were included in the analysis. All scans were acquired on a 1.5 T GE scanner (Signa HDxt, GE Healthcare, United States). Each subject underwent three scans in orthogonal planes using the single-shot Fast Spin-Echo (ssFSE) sequence. The imaging parameters were as follows: TR = 2000 ms/1041 ms, TE = 165 ms, no slice gap, in-plane resolution = 1.64×1.64 mm^2^/1.56×1.56 mm^2^, slice thickness = 4 mm.

#### Dataset 4 – Women’s Hospital of Zhejiang University Medical School (WHZ)

17 healthy pregnant women were scanned at Women’s Hospital (WHZ) from November 2018 to May 2019. After quality control, 13 fetuses at 23w to 37w GA scanned on a 1.5 T GE scanner (Signa HDxt, GE Healthcare, United States) were used for the following analysis. ssFSE data were acquired with the following imaging parameters: TR = 2400 ms, TE = 125-135 ms, no slice gap, in-plane resolution = 0.74×0.74 mm^2^/0.731×0.731 mm^2^, slice thickness = 2-5 mm.

#### Dataset 5 – Sir Run Run Shaw Hospital (RRS)

MRI scans were performed on 68 healthy pregnant women at Sir Run Run Shaw Hospital (RRS) using a 1.5 T Siemens scanner (MAGNETOM Avanto, Siemens Healthcare, Germany) from March 2020 to March 2022. After exclusions, 35 healthy singleton fetuses with GA ranging from 22 weeks to 38 weeks were included for further analysis. T2-HASTE imaging sequence was used with the following parameters: TR = 1400ms, TE = 114ms, no slice gap, in-plane resolution = 1.02×1.02 mm^2^, slice thickness = 4 mm.

### 2.2 Image processing

We utilized a standardized processing pipeline for all datasets, as illustrated in Figure 1. The orthogonal 2D multislice stacks underwent a preprocessing reconstruction pipeline using NiftyMIC (https://github.com/gift-surg/NiftyMIC) to obtain high-resolution 1 mm isotropic 3D volumes. The pipeline involved stack selection, brain extraction, bias field correction, slice-to-volume registration (SVR), and super-resolution reconstruction (SRR) [3] (Figure 1a).

**Figure 1.**
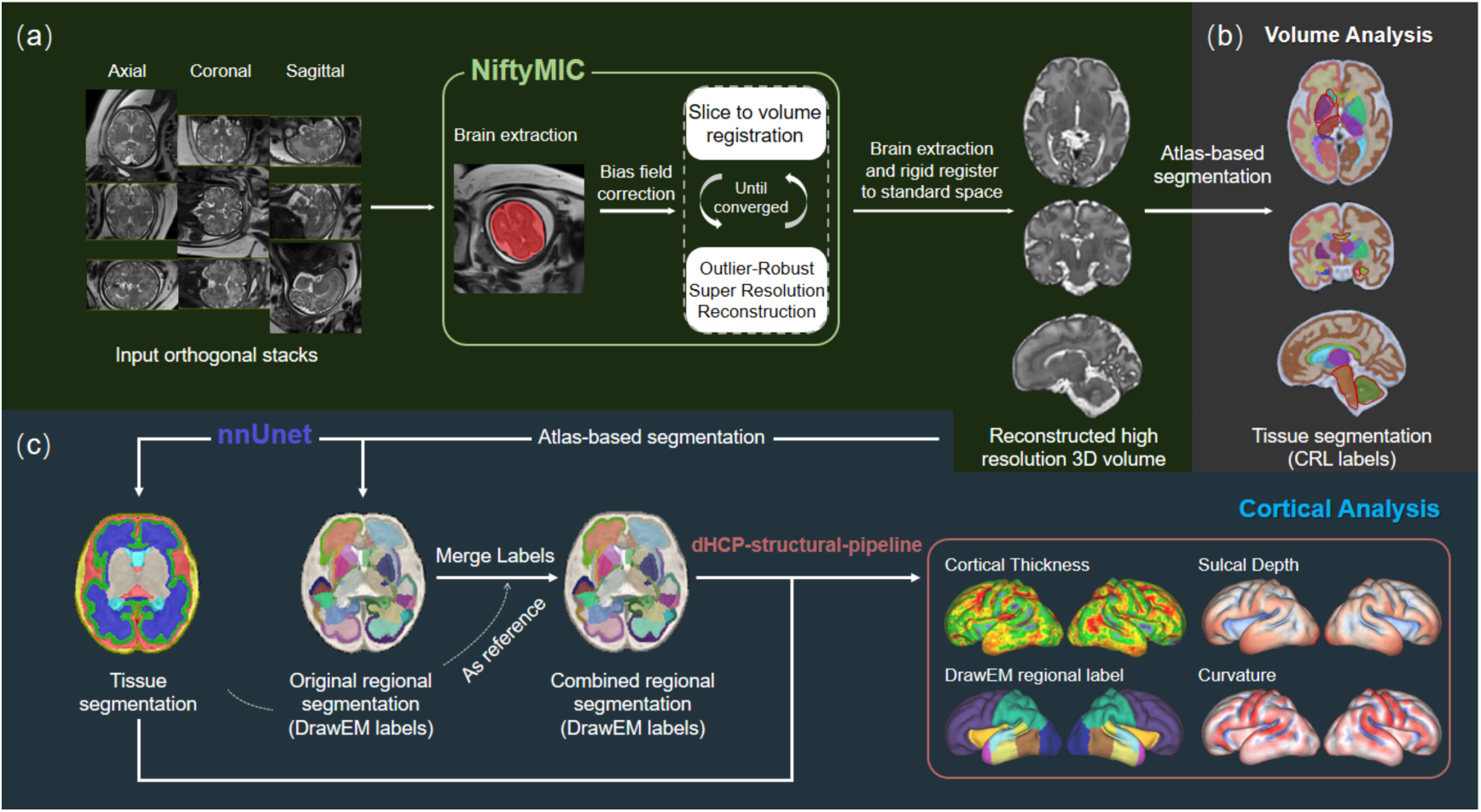
Overview of data processing pipeline including fetal brain MRI reconstruction (a) and quantification of regional brain volume (b) and cortical measurements (c).

For the extraction of deep brain structures, we employed an atlas-based segmentation method using a three-stage registration (rigid, affine, and SyN) in ANTs[36](Figure 1b). In brief, the fetal brain images were non-linearly registered to the standard CRL atlas[37] at corresponding GAs, and the segmentations in the subject space were obtained by applying the inverse transformations to the reference segmentations in CRL space. Here we focused on the volumes of the hippocampus, amygdala, thalamus, caudate, putamen, cerebellum, internal capsule, corpus callosum, and midbrain.

The cortical grey matter (cGM) segmentation was obtained using nnU-net in this study[38], which used an encoder-decoder design to learn hierarchical representations and generate corresponding segmentation maps of input volumes[39, 40]. Specifically, a 3D U-Net network was trained with a batch size of 3, patch size of 112×128×112, depth of 4, and a base channel number of 32. We used PyTorch 1.8.1 and Nvidia GeForce RTX3090 for training the model. The model was optimized using a loss function based on dice coefficient and cross-entropy, and five-fold cross-validation was employed during training. We combined nnU-net results with DrawEM-derived labels[41] that parcellate the cortex into 16 region-of-interest (ROIs) per hemisphere to obtain regional cGM segmentation. We then merged them into 6 major regions (Frontal, Parietal, Temporal, Occipital lobes, Cingulate gyrus, and Insula). Cortical morphological measurements were then computed based on the regional cGM segmentation to obtain cortical thickness, curvature, and sulcal depth in each cortical region, using dHCP-structural-pipeline[42] (originally designed for neonates), which was modified to adapt to our fetal data (Figure 1c), similar to [43].

All data were visually inspected for quality assurance before analyses by experienced radiologists (S.C., Y.H., Y.G.H, Z.Q.Q), but no manual correction was performed.

### 2.3 Atlas construction

To compare the effect of field strength and slice thickness for fetal atlas generation and also serve for different clinical application scenarios, we constructed three sets of spatiotemporal fetal brain T2 atlases based on the data from this study. Specifically, we used 96 reconstructed 3D T2w images of fetal brains scanned on 1.5 T scanners from three sites (DWC, WHZ, and RSS) to construct a *1.5T atlas.* Additionally, we utilized 122 fetal brain images acquired at SPH on a 3T scanner with 2mm and 4mm slice thickness to generate a *3T 2mm atlas* and *3T 4mm* atlas, respectively. We employed the same atlas construction pipeline as in [43].

### 2.4 Statistical analysis

In order to identify site effects of field strength (FS), manufacture (M), in-plane resolution (R), and slice thickness (ST) on subcortical volume and cortical measurements, we applied generalized additive models (GAMs) independently for each ROI. The GAMs were implemented using restricted maximum likelihood with the GA as a potential nonlinear effect (REML, implemented in mgcv package[44]) in R (https://www.r-pro-ject.org/). Each site effect was modeled with two categorical variables: FS = 3T or 1.5T, M = Siemens or GE, ST = 2 mm or 4 mm, and R = 1 for in-plane resolution higher than 1.5×1.5 mm^2^ or 0 otherwise. The GAM model was defined as follows:

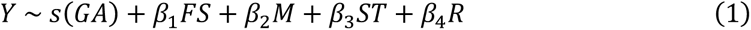

where Y indicates the morphological measurements, *s*(*GA*) represents the nonlinear effects, *β_1_, β_2_, β_3_,* and *β_4_* are the coefficients for effects of field strength, manufacture, slice thickness, and in-plane resolution. For each site effect, we used the false discovery rate (FDR) approach[45] to correct for multiple comparisons across all ROIs.

For the removal of the site effects, we applied ComBat-GAM harmonization procedure[34] to capture the age-related nonlinear change, and GA was used as the biological variable as well as the only smooth term in the model (https://github.com/rpomponio/neuroHarmonize). The harmonized data underwent GAM analysis again to evaluate the performance of ComBat-GAM. Furthermore, we investigated whether these detected site effects would affect developmental trajectories of morphological metrics. First, we quantified the developmental trajectories of subcortical volume, cortical thickness, curvature, and sulcal depth, using different growth models according to the goodness of fit, similar to [43]. Specifically, exponential growth model was chosen to fit the subcortical volume of each ROI as follows:

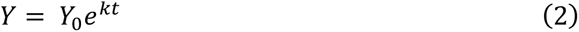

where *Y* represents the subcortical volume, *t* is the GA, k is the growth rate and *Y*_0_ is the intercept. Cortical thickness data were fitted using the Beta growth and decay model[46]. The model is defined as follows:

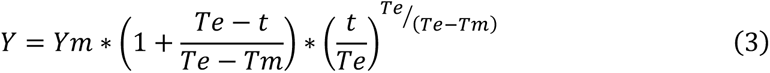

where *Ym* represents the peak cortical thickness and *Te* is the GA when cortical thickness reaches the peak, and *t* is the GA. For curvature, we used Gompertz growth model as follows:

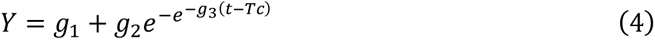

where *g*_3_ controls the growth rate and *Tc* determines the GA when peak growth of curvature occurs. The sulcal depth was fitted using the linear model:

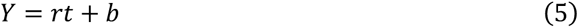

where *r* indicates the growth rate.

After fitting the models, we performed Welch’s test with Bonferroni correction to analyze statistically significant differences in the fitted parameters between groups based on the targeted site effects in each ROI (e.g., high field strength vs. low field strength), given unequal sample sizes and data variance. It’s important to note that we controlled for other site effects by removing their influence based on the GAM model results outlined in Equation (1). Further grouping details can be found in Table S1.

## 3 Results

### 3.1 Visualization and identification of site effects

#### Visual Comparison

Visual inspection of the reconstructed 3D T2-weighted volumes of the fetal brains at 32-33 weeks of GA from the five datasets indicated that the 3T data (SPH) and higher in-plane resolution data (WHZ) exhibited higher signal-to-noise ratio (SNR) compared to the 1.5T data and lower in-plane resolution data at the individual level (Figure 2). The corresponding subcortical segmentation by atlas-based segmentation method and cGM segmentation by nnU-net showed consistent fetal brain segmentation across all five datasets (Figure 2).

**Figure 2.**
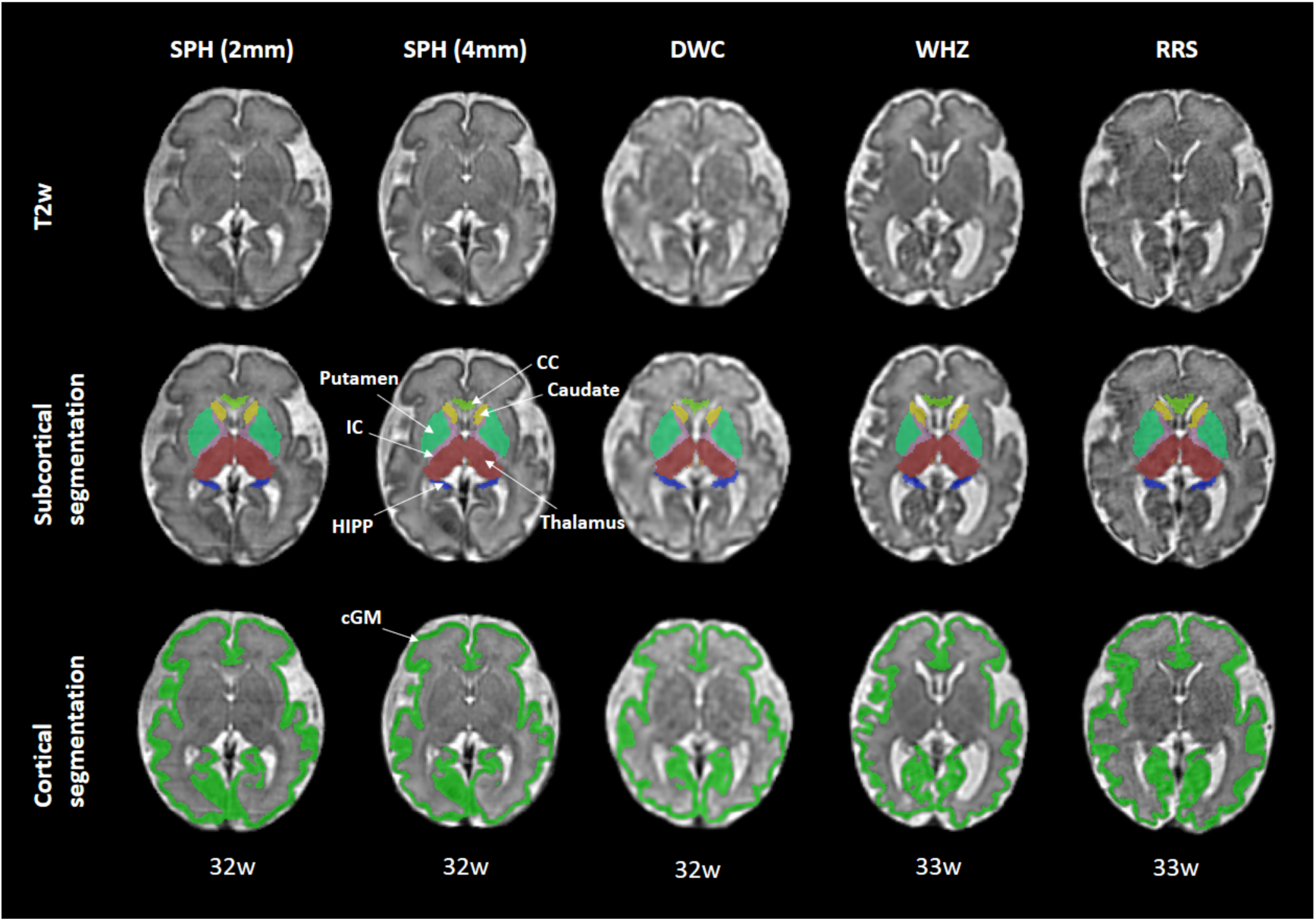
3D T2w images of fetal brains at 32-33 weeks of GA and corresponding subcortical segmentation and cortical gray matter segmentation results from 5 datasets for visual comparison. Abbreviations: CC = corpus callosum, IC = internal capsule, HIPP = hippocampus, cGM = cortical gray matter. SPH (2mm) stands for SPH data acquired at 2mm slice thickness while SPH (4mm) stands for those at 4mm slice thickness.

#### Effects of field strength and slice thickness on subcortical volume measurements

The influence of each site factor on volume of each subcortical ROI is summarized in Table S2, and only two subcortical ROIs showed significant site effects. The GAM results revealed significantly lower volume measurements of the corpus callosum for 3T data compared to 1.5T (left panel in Figure 3a). Additionally, the volume of the internal capsule was significantly impacted by slice thickness, with larger volumes in data acquired with a 4 mm slice thickness compared to a 2 mm slice thickness (right panel in Figure 3a). No detectable influences on subcortical regional volumes were found for manufacture or in-plane resolution. After ComBat-GAM harmonization, the significant effects of field strength and slice thickness on volume measurements were successfully eliminated, while preserving the developmental trend of subcortical regional volumes (Table S2 and Figure 3b).

**Figure 3.**
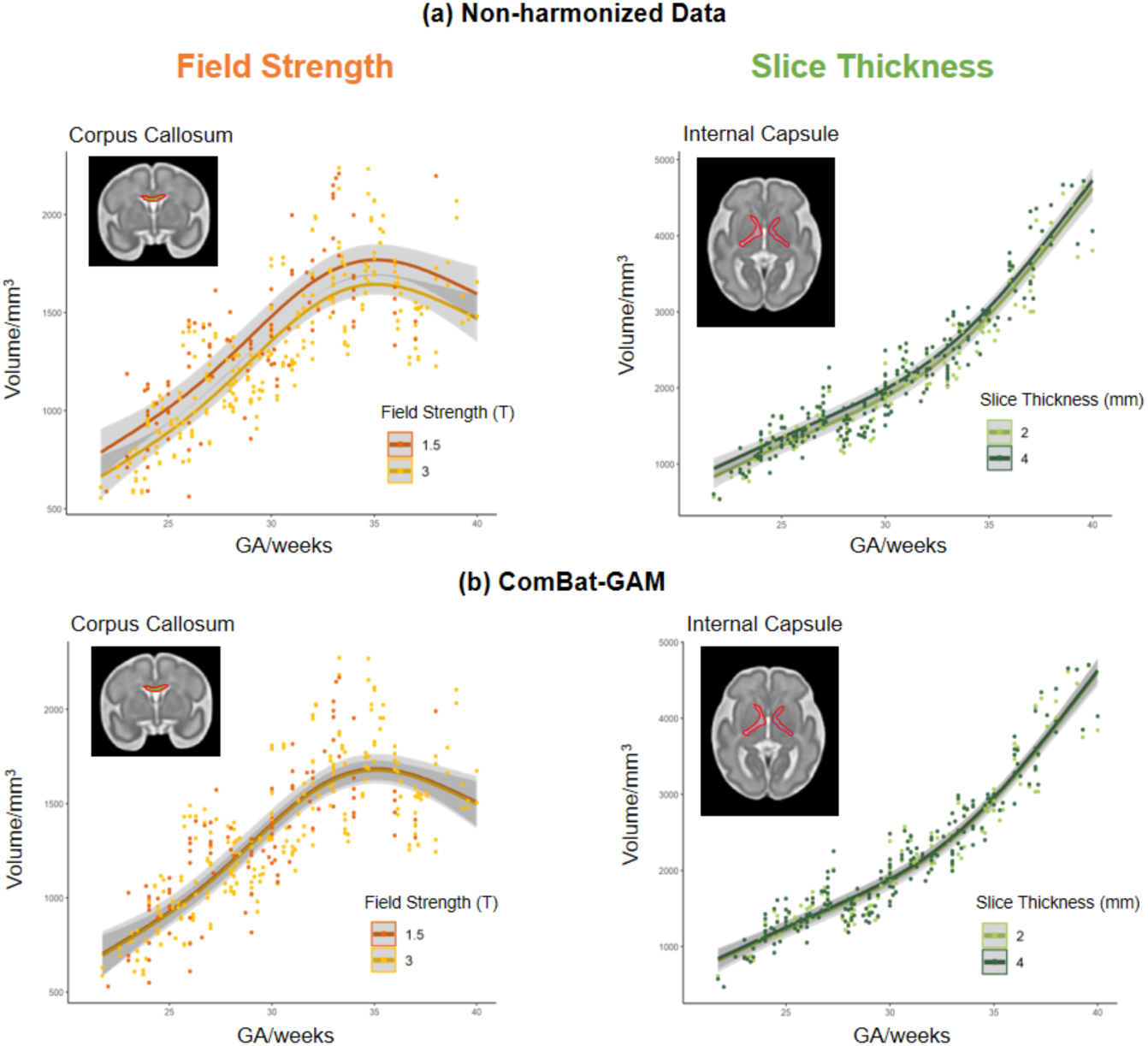
Effects of field strength and slice thickness on volume measurements in corpus callosum and internal capsule. Age-related changes in non-harmonized volume data (a) and harmonized volume data (b) using different field strengths and slice thicknesses.

#### Effects of field strength, in-plane resolution, and manufacture on cortical measurements

In general, GAM analysis detected significant effects of field strength, in-plane resolution, and manufacture on the three cortical morphological measurements in multiple regions (Table S3-S5). Figure 4 displays representative ROIs where the site effects were the most significant. Specifically, we found significantly lower cortical thickness, sulcal depth, and curvature for 3T data compared to 1.5T scanners (Figure 4b). Curvature and sulcal depth calculated from GE scanner data showed smaller values in comparison to those acquired from Siemens scanners (Figure 4c). In-plane resolution showed the opposite effect on cortical thickness versus sulcal depth, i.e., higher in-plane resolution resulted in lower cortical thickness but higher sulcal depth (Figure 4d). The site effects on cortical morphology were visualized in Figure 5 for all ROIs. Note that the sulcal depth of insula and cingulate gyrus exhibited different responses to the site effects.

**Figure 4.**
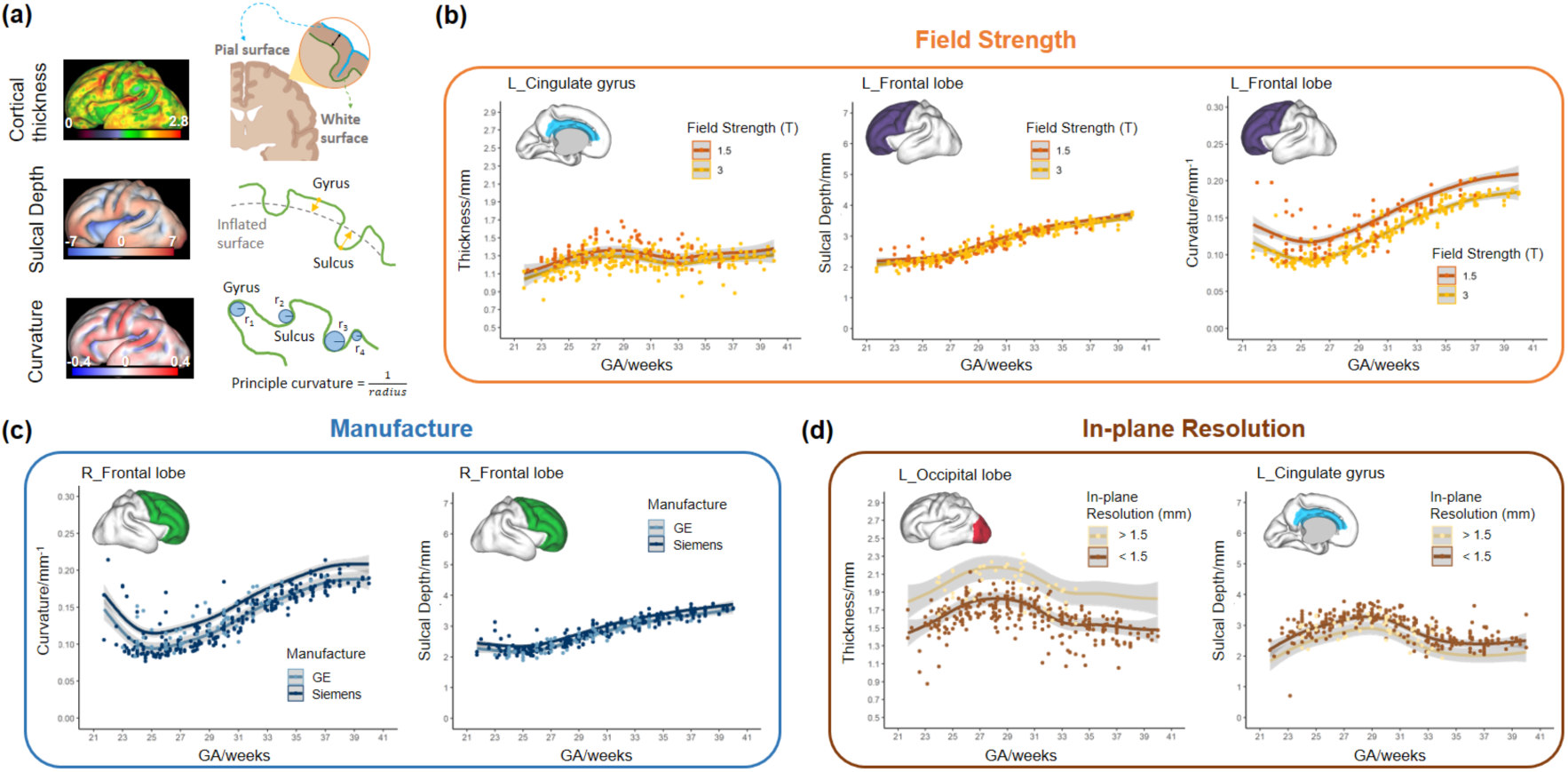
(a) Graphical illustrations of cortical thickness, sulcal depth, and curvature measurements. Effects of field strength (b, 1.5T in orange and 3T in yellow), manufacture (c, GE scanner in light blue and Siemens in darker blue) and in-plane resolution (d, higher resolution in brown and lower in beige) on cortical morphological measurements in representative ROIs.

**Figure 5.**
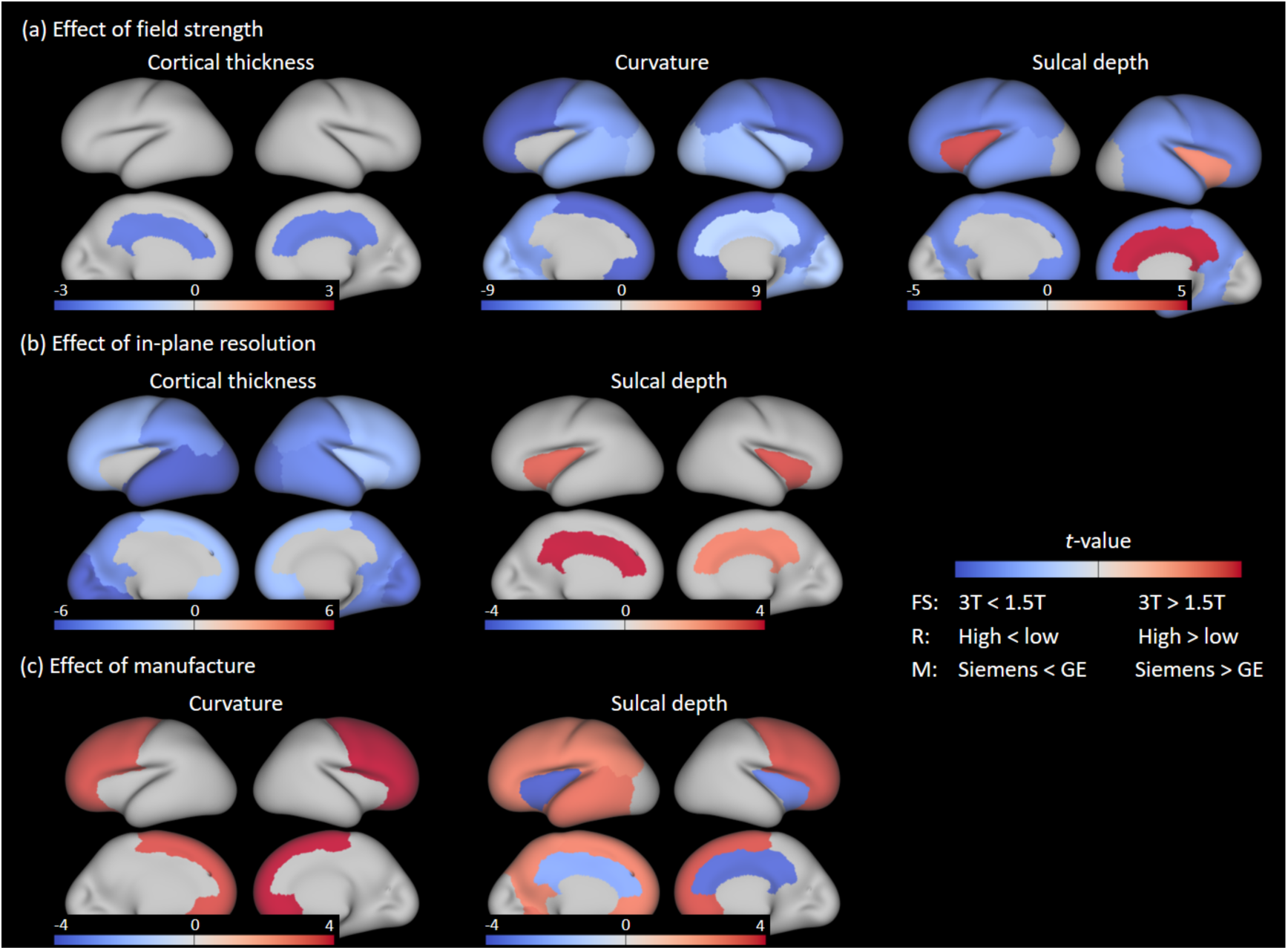
t-value map of site effects on cortical morphological measurements in regions with significant effects. Abbreviations: FS = Field Strength, R = in-plane Resolution, M = Manufacture.

ComBat-GAM effectively removed the site effects on cortical morphological measurements (Table S3-S5). Assuming T2w data scanned with higher field strength, higher in-plane resolution, and thinner slice thickness gives more accurate morphological measurements due to better SNR, and lower field strength and lower resolution generally lead to overestimation of subcortical volume and cortical morphological measurements. Regarding manufacture effect, curvature and sulcal depth of most regions showed larger values in Siemens data.

**Table 2.** Summarized site effects on subcortical volume and cortical morphological measurements.

|  | Field Strength | Manufacture | Slice Thickness | In-plane Resolution |
| --- | --- | --- | --- | --- |
| Volume | 3T < 1.5T |  | 2mm < 4mm |  |
| Thickness | 3T < 1.5T |  |  | High < Low |
| Curvature | 3T < 1.5T | GE < Siemens |  |  |
| Sulcal Depth | 3T < 1.5T* | GE < Siemens* |  | High > Low<sup>#</sup> |

Group1 < Group 2: The value of the morphological index is smaller in Group 1 than that in Group 2
\* Insula and cingulate gyrus showed opposite site effects trend to other regions
<sup>#</sup> Only the insula and cingulate gyrus had significant in-plane resolution effects

### 3.2 Influence of site effects on developmental trajectory analysis

We portrayed developmental trajectories of the fetal brains using previously defined growth models and analyzed whether the detected site effects from section 3.1 influenced the developmental patterns. We detected that only the parameter *Ym* (peak cortical thickness) in cortical thickness trajectories was significantly influenced by the effects of field strength and in-plane resolution (Table S6). The peak cortical thickness (*Ym*) fitted from data with low field strength or low in-plane resolution was significantly larger compared to those from higher field strength or higher in-plane resolution data (Figure 6), consistent with the findings in 3.1. These significant differences were efficiently mitigated through ComBat-GAM harmonization process. Results in this section indicated that the temporal developmental pattern is relatively robust to site effects.

**Figure 6.**
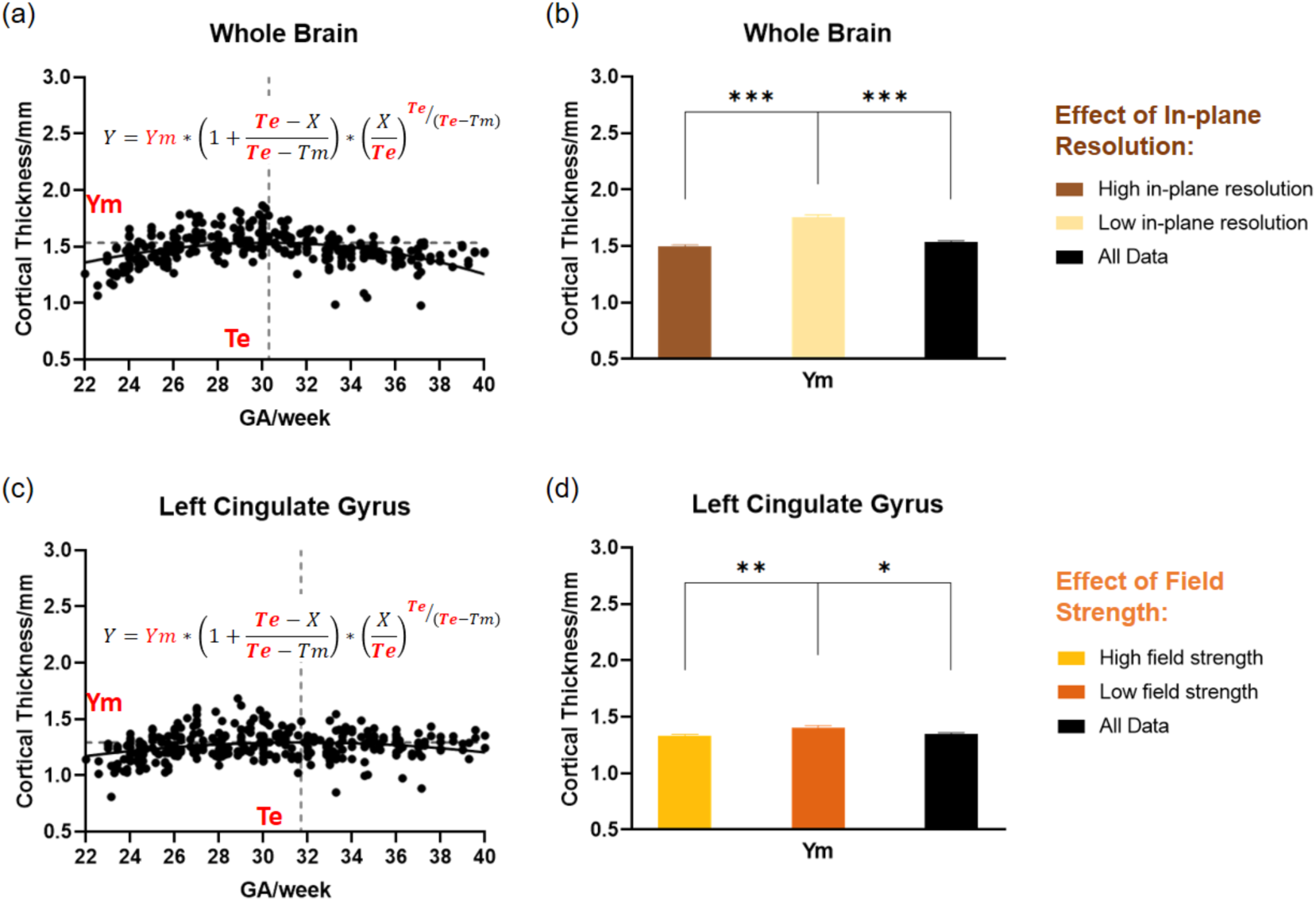
In-plane resolution and field strength effects on *Ym* of Beta Growth then Decay model of cortical thickness in representative regions. Developmental trajectories of cortical thickness across GA in whole brain (a) and left cingulate gyrus (c). Fitted *Ym* of cortical thickness trajectory in whole brain (b) and left cingulate gyrus (d), using data acquired at different in-plane resolution (higher resolution in brown and lower in beige) and different field strength (higher field strength in yellow and lower in orange). * Adjusted P < 0.05, ** adjusted P < 0.01, and *** adjusted P < 0.001.

### 3.3 Fetal brain atlases generated for different site effects

We generated three sets of spatiotemporal fetal brain T2 atlases (1.5T 4mm thickness, 3T 4mm thickness, and 3T 2mm thickness) to check whether the site effects were visible in the group averaged atlas level. These atlases also provided reference templates for different purposes, e.g., studies using clinical data (4mm thickness) versus research data (2mm thickness) or low field versus high field applications. Specifically, we constructed a *1.5T atlas* of fetal brains at 23w to 36w GA and a *3T 4mm and 3T 2mm atlas* of the fetal brains at 23w to 38w GA. Upon visual comparison (Figure 7), we observed that *1.5T atlas* had slightly lower SNR compared to the other two *3T* atlases, but the quality was still satisfying. The brain structures and brain sizes were generally consistent between these three versions of the atlases. These atlases can be found at the following GitHub repository: https://github.com/zjuwulab/Multisite-fetal-brain-atlas. Note the *3T 2mm atlas* is the same as previously published fetal brain atlas[43].

**Figure 7.**
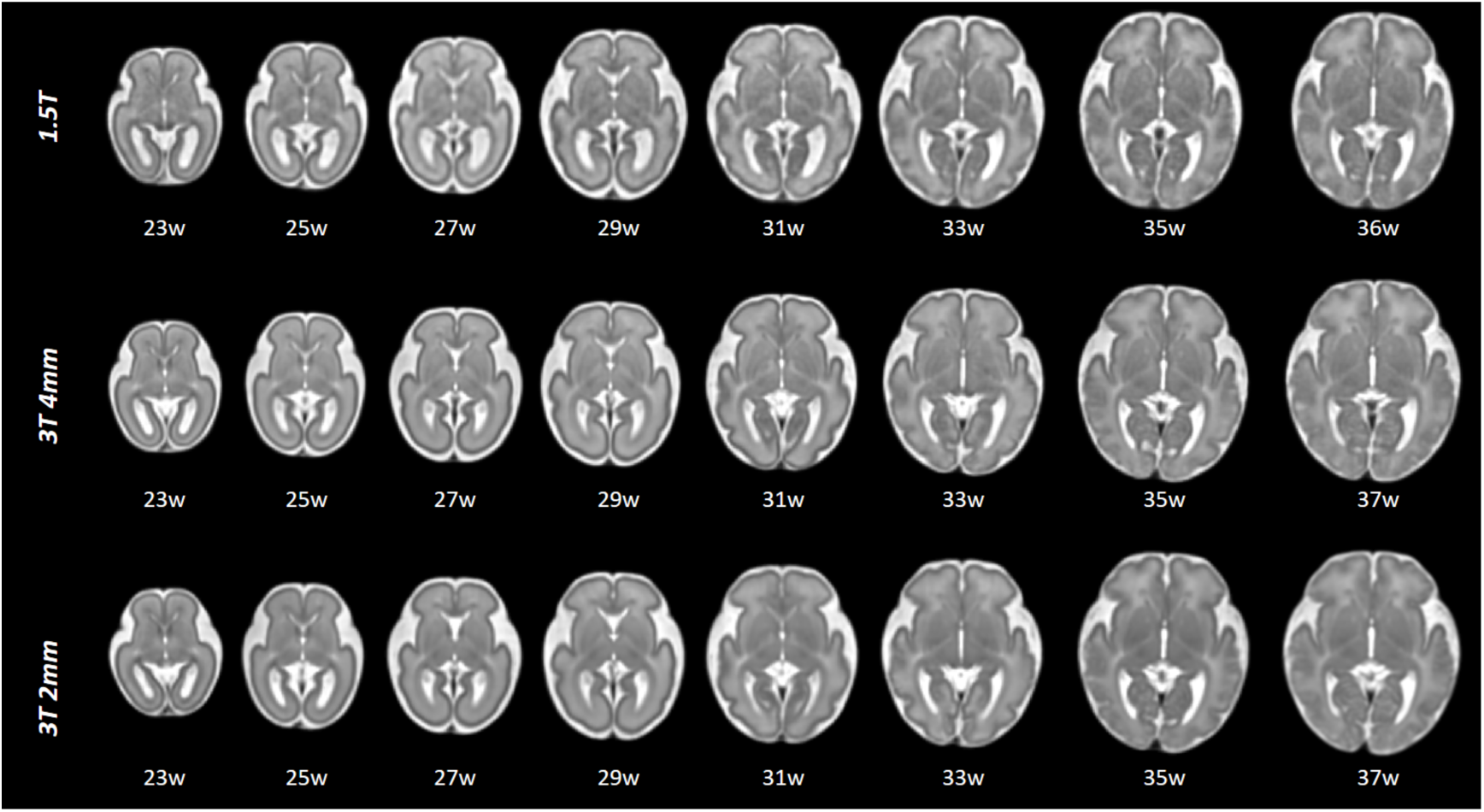
Spatiotemporal fetal brain atlases constructed based on data from 1.5 T field strength (top, 1.5T atlas), 3 T field strength with 4 mm (middle) and 2mm (bottom) slice thickness.

## 4 Discussion

In this study, we systematically investigated the multisite effects on fetal brain MRI, with a focus on brain morphological measurements during early development. Our findings provide important guidelines for the fetal MRI community when scanning the fetal brains, e.g., low field strength and low resolution may lead to overestimation of brain volume and cortical measurements and these effects were region-specific. Additionally, we showed the effectiveness of the ComBat-GAM method in removing these site effects while preserving the developmental trend. Furthermore, we generated spatiotemporal fetal brain atlases for different field strengths and imaging resolutions, which could serve diverse clinical requirements.

We identified field strength contributed significantly to all four studied morphological measurements: subcortical volume, cortical thickness, curvature, and sulcal depth. Our results demonstrated higher estimates of subcortical volume and cortical measurements in 1.5T data compared to 3T data. There is significant literature documenting the field strength effect on regional volume[47] and cortical thickness[18, 19, 21, 22] in adults and neonates which showed a decrease of these measurements on 1.5T than 3T, based on T1-weighted MRI. Han et al. pointed out that the effect that mean cortical thickness values are higher at 3T than at 1.5T is largely due to the field strength affecting the longitudinal relaxation time T1, which changes the intensity and contrast of the images[19]. Therefore, we hypothesized that the opposite field strength effect detected on fetal morphological measurements compared to adults and neonates may be attributed to the differences in imaging modalities, that the 3D structural MRI used for adult and neonatal studies mentioned above are T1w images, but fetal brain studies used T2w images as it gives better contrast than T1w during fetal stage. Also, note that the image intensity contrast between cortical gray matter and white matter in the fetal brains is opposite compared to the adult brain.

We detected significant scanner manufacture effects on curvature and sulcal depth measurements, with lower values observed on GE scanners compared to Siemens scanners in most regions (except for insula and cingulate). Previous studies have reported that manufacture effect was detectable in morphological structural studies results[17, 18, 48–51]. However, the manufacture effect on volumetric and cortical measurements varied in different studies. Reig et al. reported that Siemens tended to show higher white matter volumes compared with GE or Philips[17]. Fortin et al. showed there was a higher cortical thickness value on GE scanners than on Siemens scanners in adult brains [18]. On the contrary, Fennema et al. showed the estimated total intracranial volume (eTIV) was higher on Siemens than that from GE[51]. We think the manufacture effect is difficult to interpret because it is often coupled with other imaging parameters and hardware setup (coil sensitivity, field inhomogeneity, etc.), and protocol differences on different scanner prototypes. Fortunately, the manufacture effect on fetal brain MRI can be effectively removed after harmonization.

In terms of the regional difference in response to site effect, we observe that the sulcal depth of insula and cingulate gyrus consistently showed opposite trends compared to other regions and metrics. This may be due to the different developmental patterns of sulcal depth in these regions with GA, which exhibited complex age-related nonlinear patterns in the insula and cingulate gyrus compared to the monotonically increasing developmental trends observed in other brain regions (see supplementary Figure S2).

To address the site effect, a number of harmonization methods were proposed to minimize or even eliminate batch effects in neuroimaging studies. In general, existing harmonization methods could be categorized into statistical harmonization and deep learning based harmonization which can be further subdivided based on whether the methods are designed for retrospective or prospective study[52]. In this study, we employed ComBat-GAM[34] which is a retrospective statistical harmonization method. Sun et al. performed a comparison study of four harmonization methods on cortical thickness across lifespan and found that ComBat-GAM provided a stronger estimate of age-related nonlinear thickness reduction, compared to ComBat and linear mixed effects models[53]. In our study, most morphological measurements exhibited nonlinear changes across GA. Therefore, we chose ComBat-GAM to harmonize our data, which successfully removed site effects while preserving age-related effects.

Overall, our results suggested that high field strength and high in-plane resolution are preferred for analysis of fetal brain morphology, and harmonization procedures are necessary for multisite studies.

### 4.1 Limitation

Despite the valuable insights gained from this study, there are several limitations that should be considered. First, due to the challenges associated with collecting fetal data from the same subjects across different sites, we were unable to perform a paired comparison or perform a travel phantom experiment. Second, the sample size in our study, although relatively large for fetal brain imaging, was still relatively small and the subjects were not equally distributed across sites. Larger and more homogeneous datasets would provide higher statistical power and enhance the generalizability of our findings. Lastly, our study focused on four specific morphological indicators, and the site effects investigated were primarily limited to field strength, manufacturer, in-plane resolution, and slice thickness. Future studies could include additional morphological indicators and other effects that may affect the fetal MRI findings, such as image preprocessing methods.

## 5 Conclusion

Our study systematically investigated the effects of manufacture, field strength, in-plane resolution, and slice thickness on the subcortical volumes and cortical measurements in healthy fetuses across multiple centers. Our findings highlighted the significant influence of field strength and slice thickness on volumes of only two subcortical structures, while cortical morphology was strongly affected by field strength, manufacture, and in-plane resolution, with lower field strength and imaging resolution leading to overestimation of these measurements. Moreover, the developmental trajectories of cortical thickness were found to be related to field strength and in-plane resolution. Additionally, we demonstrated that ComBat-GAM can effectively remove nuisance variabilities associated with the acquisition while maintaining developmental changes. Therefore, it is feasible to pool multicenter fetal brain MRI for studying normal brain development and fetal brain diseases.

## Supplementary material

**Table S1.** Grouping strategy of five fetal brain MRI datasets for targeted site effects on developmental trajectories analysis. Note that ‘D’ stands for Dataset.

|  | Field Strength | Manufacture | Slice Thickness | In-plane Resolution |
| --- | --- | --- | --- | --- |
| Group1 | 3T (D1, D2) | GE (D3-4) | 2mm (D1) | High (D1-2, D4-5) |
| Group2 | 1.5T (D3-5) | Siemens (D1-2, D5) | 4mm (D2-5) | Low (D3) |

**Table S2.** Summary of site-related effects (β values) on subcortical regional volumes before and after harmonization.

| *Subcortical Volumes* | Non-harmonized raw data |  |  |  | After ComBat-GAM harmonization |  |  |  |
| --- | --- | --- | --- | --- | --- | --- | --- | --- |
| | Field Strength ($\beta_1$) | Manufacture ($\beta_2$) | Slice Thickness ($\beta_3$) | In-plane Resolution ($\beta_4$) | Field Strength ($\beta_1$) | Manufacture ($\beta_2$) | Slice Thickness ($\beta_3$) | In-plane Resolution ($\beta_4$) |
| Hippocampus | -49.995 | 16.934 | 4.344 | 37.028 | 11.749 | -16.821 | -7.750 | 3.777 |
| Amygdala | -4.559 | 12.420 | 2.887 | -9.551 | 3.851 | 4.064 | -1.119 | -13.002 |
| Thalamus | -134.625 | 145.625 | 67.076 | 68.592 | -9.218 | 29.922 | 6.617 | -5.429 |
| Caudate | -85.213 | 60.163 | 23.103 | -14.212 | -3.057 | 7.592 | 0.388 | -10.102 |
| Putamen | -77.043 | 311.890 | 117.823 | -171.872 | 17.755 | 92.499 | 23.650 | -106.008 |
| Cerebellum | -163.562 | 495.186 | 134.639 | -6.847 | 114.724 | 23.771 | -19.661 | -47.160 |
| Internal Capsule | -60.626 | 40.156 | 105.866** | 75.917 | 7.188 | -29.072 | 29.196 | 30.582 |
| Corpus Callosum | -124.659* | 4.003 | -27.207 | 28.888 | -8.467 | -38.983 | -15.639 | 26.694 |
| Midbrain | -176.740 | 240.286 | 25.168 | -12.020 | -7.093 | 48.609 | -13.624 | -19.550 |

\* Adjusted $P < 0.05$ and \*\* adjusted $P < 0.01$

**Table S3.**
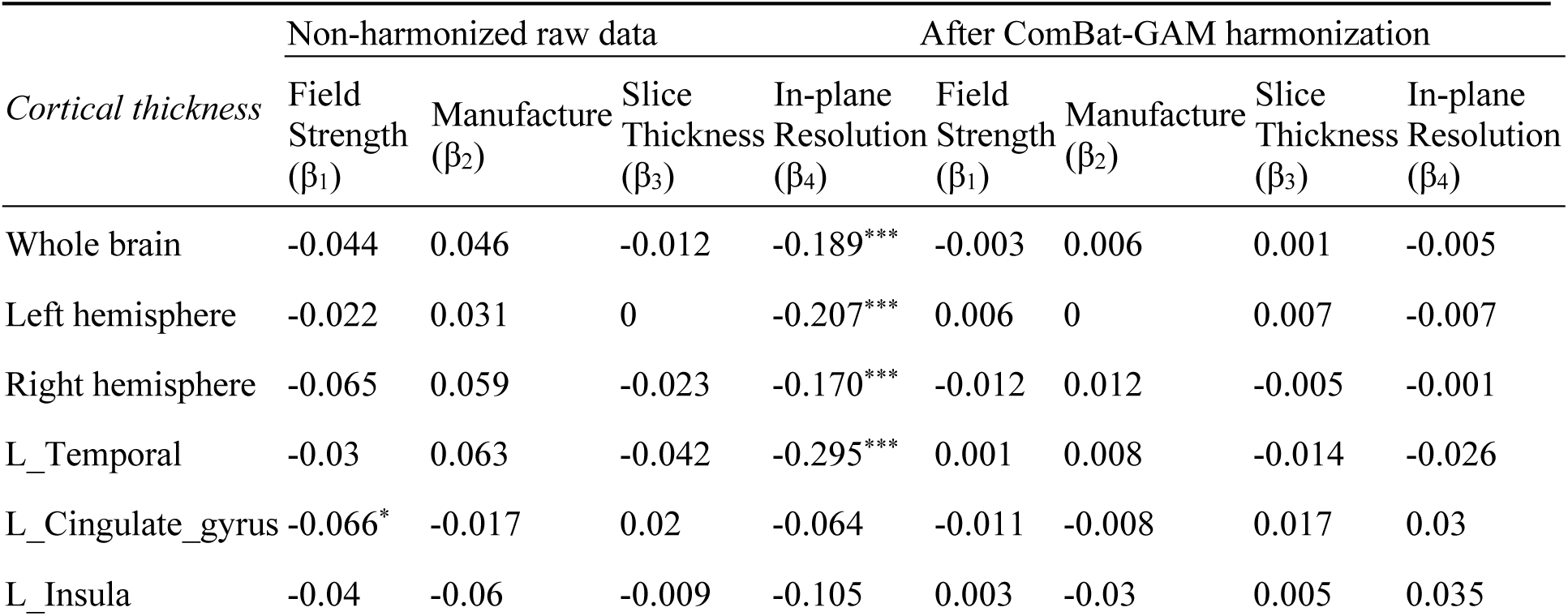
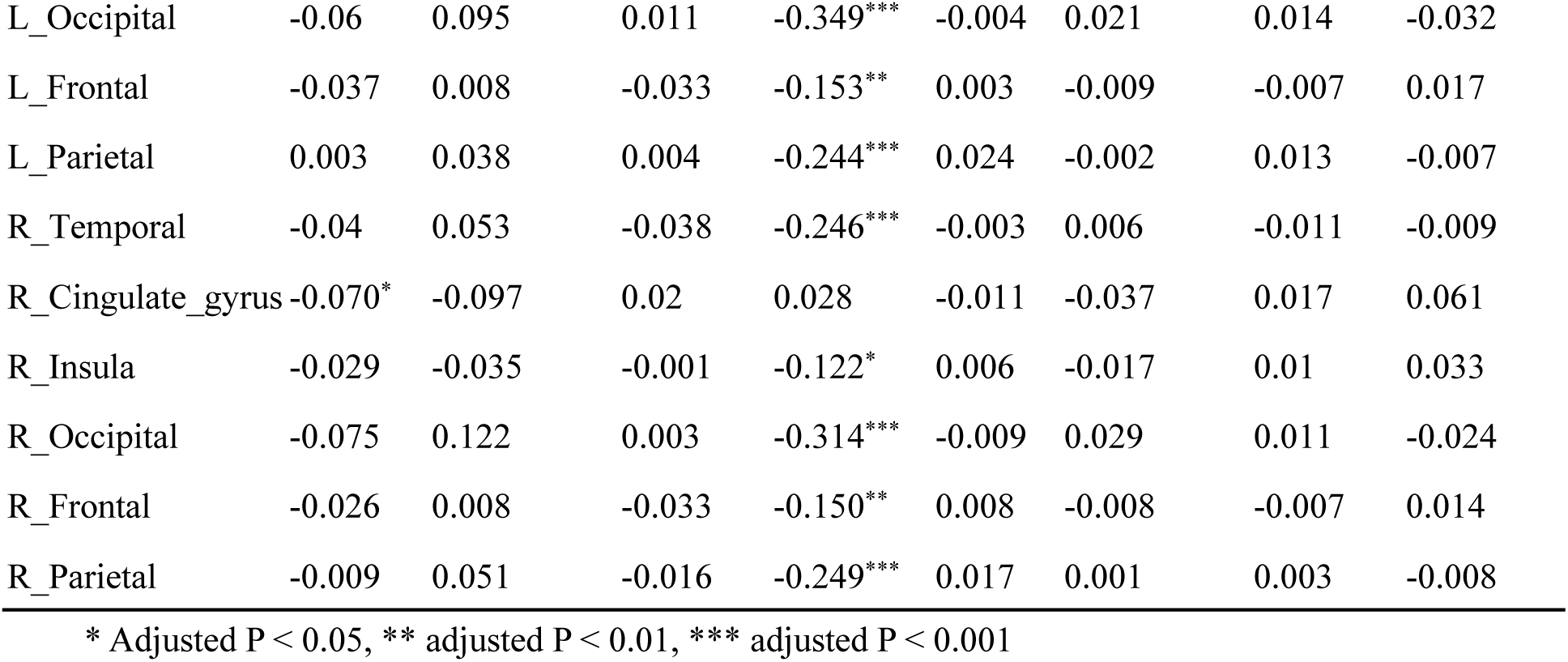
Summary of site-related effects (β values) on cortical thickness before and after harmonization.

| *Cortical thickness* | Non-harmonized raw data |  |  |  | After ComBat-GAM harmonization |  |  |  |
| --- | --- | --- | --- | --- | --- | --- | --- | --- |
| | Field Strength ($\beta_1$) | Manufacture ($\beta_2$) | Slice Thickness ($\beta_3$) | In-plane Resolution ($\beta_4$) | Field Strength ($\beta_1$) | Manufacture ($\beta_2$) | Slice Thickness ($\beta_3$) | In-plane Resolution ($\beta_4$) |
| Whole brain | -0.044 | 0.046 | -0.012 | -0.189*** | -0.003 | 0.006 | 0.001 | -0.005 |
| Left hemisphere | -0.022 | 0.031 | 0 | -0.207*** | 0.006 | 0 | 0.007 | -0.007 |
| Right hemisphere | -0.065 | 0.059 | -0.023 | -0.170*** | -0.012 | 0.012 | -0.005 | -0.001 |
| L_Temporal | -0.03 | 0.063 | -0.042 | -0.295*** | 0.001 | 0.008 | -0.014 | -0.026 |
| L_Cingulate_gyrus | -0.066* | -0.017 | 0.02 | -0.064 | -0.011 | -0.008 | 0.017 | 0.03 |
| L_Insula | -0.04 | -0.06 | -0.009 | -0.105 | 0.003 | -0.03 | 0.005 | 0.035 |
| L_Occipital | -0.06 | 0.095 | 0.011 | -0.349*** | -0.004 | 0.021 | 0.014 | -0.032 |
| L_Frontal | -0.037 | 0.008 | -0.033 | -0.153** | 0.003 | -0.009 | -0.007 | 0.017 |
| L_Parietal | 0.003 | 0.038 | 0.004 | -0.244*** | 0.024 | -0.002 | 0.013 | -0.007 |
| R_Temporal | -0.04 | 0.053 | -0.038 | -0.246*** | -0.003 | 0.006 | -0.011 | -0.009 |
| R_Cingulate_gyrus | -0.070* | -0.097 | 0.02 | 0.028 | -0.011 | -0.037 | 0.017 | 0.061 |
| R_Insula | -0.029 | -0.035 | -0.001 | -0.122* | 0.006 | -0.017 | 0.01 | 0.033 |
| R_Occipital | -0.075 | 0.122 | 0.003 | -0.314*** | -0.009 | 0.029 | 0.011 | -0.024 |
| R_Frontal | -0.026 | 0.008 | -0.033 | -0.150** | 0.008 | -0.008 | -0.007 | 0.014 |
| R_Parietal | -0.009 | 0.051 | -0.016 | -0.249*** | 0.017 | 0.001 | 0.003 | -0.008 |

\* Adjusted P < 0.05, \*\* adjusted P < 0.01, \*\*\* adjusted P < 0.001

**Table S4.** Summary of site-related effects (β values) on cortical curvature before and after harmonization.

| *Curvature* | Non-harmonized raw data |  |  |  | After ComBat-GAM harmonization |  |  |  |
| --- | --- | --- | --- | --- | --- | --- | --- | --- |
| | Field Strength ($\beta_1$) | Manufacture ($\beta_2$) | Slice Thickness ($\beta_3$) | In-plane Resolution ($\beta_4$) | Field Strength ($\beta_1$) | Manufacture ($\beta_2$) | Slice Thickness ($\beta_3$) | In-plane Resolution ($\beta_4$) |
| Whole brain | -0.013*** | 0.006 | 0 | -0.002 | -0.004 | 0.007 | 0.001 | -0.006 |
| Left hemisphere | -0.016*** | 0.008* | -0.002 | -0.002 | 0.006 | 0.001 | 0.007 | -0.009 |
| Right hemisphere | -0.01*** | 0.004 | 0.001 | -0.003 | -0.012 | 0.011 | -0.005 | -0.001 |
| L_Temporal | -0.011*** | 0.004 | -0.002 | 0.001 | -0.002 | 0.011 | -0.014 | -0.026 |
| L_Cingulate_gyrus | -0.005 | -0.002 | 0 | 0.004 | -0.009 | -0.012 | 0.017 | 0.032 |
| L_Insula | -0.004 | 0.003 | 0.003 | -0.003 | 0.003 | -0.03 | 0.005 | 0.035 |
| L_Occipital | -0.012*** | 0.009 | -0.003 | -0.005 | -0.003 | 0.019 | 0.014 | -0.031 |
| L_Frontal | -0.025*** | 0.016** | -0.001 | -0.007 | -0.001 | -0.004 | -0.007 | 0.017 |
| L_Parietal | -0.013*** | 0.007 | 0.002 | -0.002 | 0.024 | -0.004 | 0.013 | -0.004 |
| R_Temporal | -0.008*** | 0.001 | -0.002 | 0.002 | -0.001 | 0.005 | -0.011 | -0.012 |
| R_Cingulate_gyrus | -0.011** | 0.012 | -0.006 | 0.002 | -0.01 | -0.038 | 0.017 | 0.061 |
| R_Insula | -0.01*** | 0 | 0.001 | 0.007 | 0.006 | -0.017 | 0.01 | 0.033 |
| R_Occipital | -0.009** | 0.002 | -0.002 | -0.003 | -0.009 | 0.029 | 0.01 | -0.024 |
| R_Frontal | -0.024*** | 0.021*** | -0.001 | -0.012 | 0.006 | -0.007 | -0.007 | 0.016 |
| R_Parietal | -0.018*** | 0.008 | 0.001 | 0.002 | 0.017 | 0.001 | 0.003 | -0.009 |

\* Adjusted P < 0.05, \*\* adjusted P < 0.01, \*\*\* adjusted P < 0.001

**Table S5.**
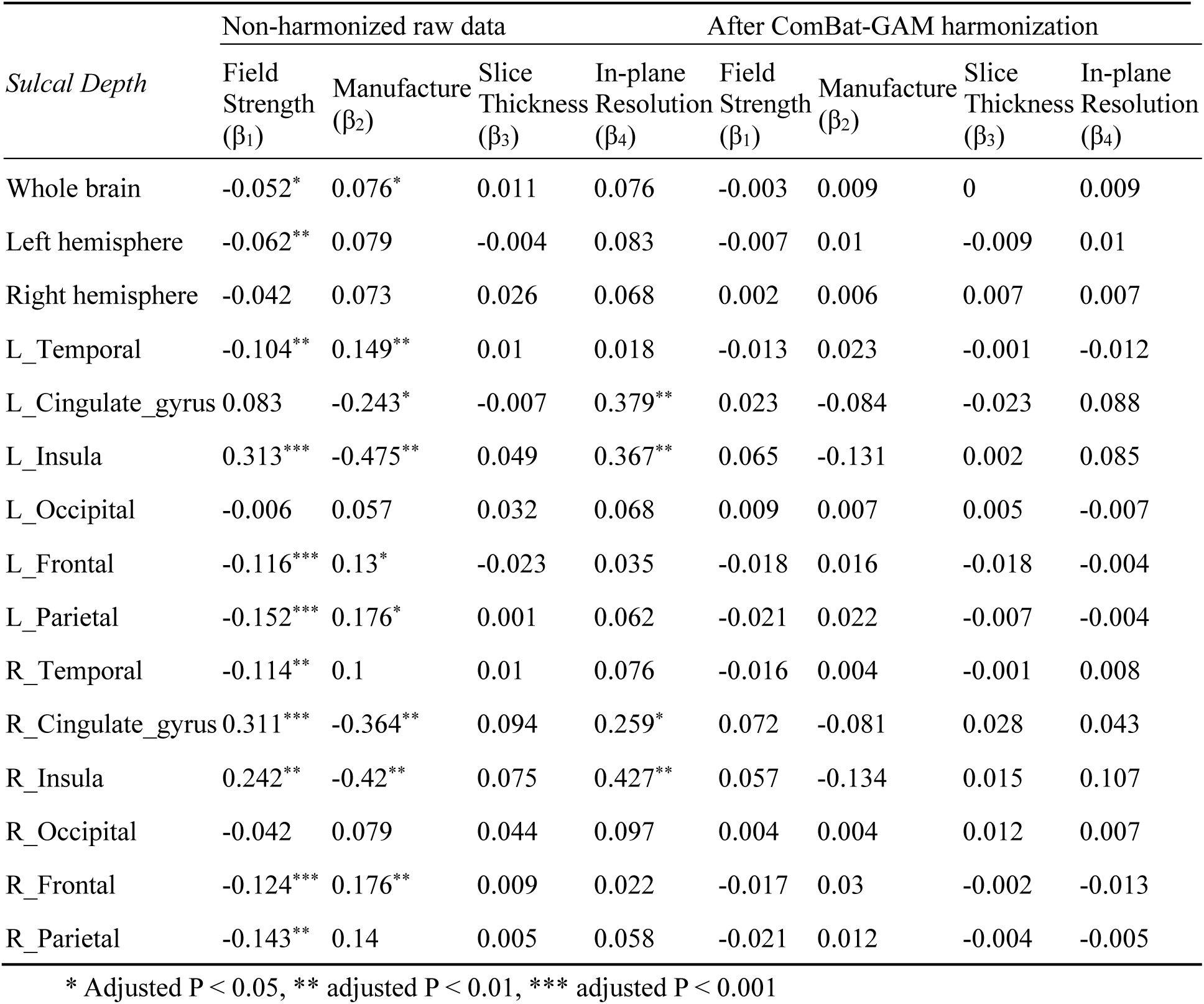
Summary of site-related effects (β values) on cortical sulcal depth before and after harmonization.

| *Sulcal Depth* | Non-harmonized raw data |  |  |  | After ComBat-GAM harmonization |  |  |  |
| --- | --- | --- | --- | --- | --- | --- | --- | --- |
| | Field Strength ($\beta_1$) | Manufacture ($\beta_2$) | Slice Thickness ($\beta_3$) | In-plane Resolution ($\beta_4$) | Field Strength ($\beta_1$) | Manufacture ($\beta_2$) | Slice Thickness ($\beta_3$) | In-plane Resolution ($\beta_4$) |
| Whole brain | -0.052* | 0.076* | 0.011 | 0.076 | -0.003 | 0.009 | 0 | 0.009 |
| Left hemisphere | -0.062** | 0.079 | -0.004 | 0.083 | -0.007 | 0.01 | -0.009 | 0.01 |
| Right hemisphere | -0.042 | 0.073 | 0.026 | 0.068 | 0.002 | 0.006 | 0.007 | 0.007 |
| L_Temporal | -0.104** | 0.149** | 0.01 | 0.018 | -0.013 | 0.023 | -0.001 | -0.012 |
| L_Cingulate_gyrus | 0.083 | -0.243* | -0.007 | 0.379** | 0.023 | -0.084 | -0.023 | 0.088 |
| L_Insula | 0.313*** | -0.475** | 0.049 | 0.367** | 0.065 | -0.131 | 0.002 | 0.085 |
| L_Occipital | -0.006 | 0.057 | 0.032 | 0.068 | 0.009 | 0.007 | 0.005 | -0.007 |
| L_Frontal | -0.116*** | 0.13* | -0.023 | 0.035 | -0.018 | 0.016 | -0.018 | -0.004 |
| L_Parietal | -0.152*** | 0.176* | 0.001 | 0.062 | -0.021 | 0.022 | -0.007 | -0.004 |
| R_Temporal | -0.114** | 0.1 | 0.01 | 0.076 | -0.016 | 0.004 | -0.001 | 0.008 |
| R_Cingulate_gyrus | 0.311*** | -0.364** | 0.094 | 0.259* | 0.072 | -0.081 | 0.028 | 0.043 |
| R_Insula | 0.242** | -0.42** | 0.075 | 0.427** | 0.057 | -0.134 | 0.015 | 0.107 |
| R_Occipital | -0.042 | 0.079 | 0.044 | 0.097 | 0.004 | 0.004 | 0.012 | 0.007 |
| R_Frontal | -0.124*** | 0.176** | 0.009 | 0.022 | -0.017 | 0.03 | -0.002 | -0.013 |
| R_Parietal | -0.143** | 0.14 | 0.005 | 0.058 | -0.021 | 0.012 | -0.004 | -0.005 |

\* Adjusted P < 0.05, \*\* adjusted P < 0.01, \*\*\* adjusted P < 0.001

**Table S6.**
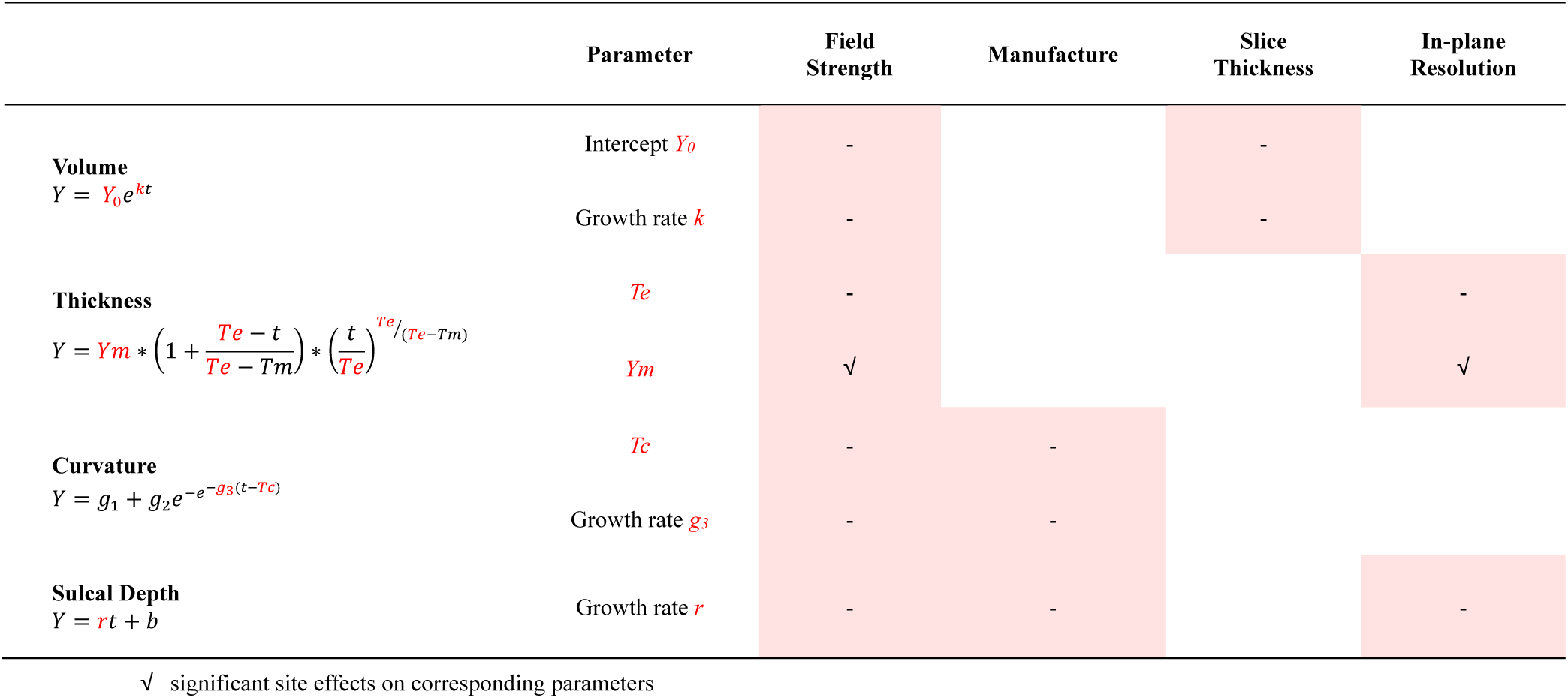
Influence of site effects detected from section 3.1 (red-filled blocks) for volume, cortical thickness, curvature, and sulcal depth on developmental trajectories parameters of different morphological measurements.

**Figure S1.**
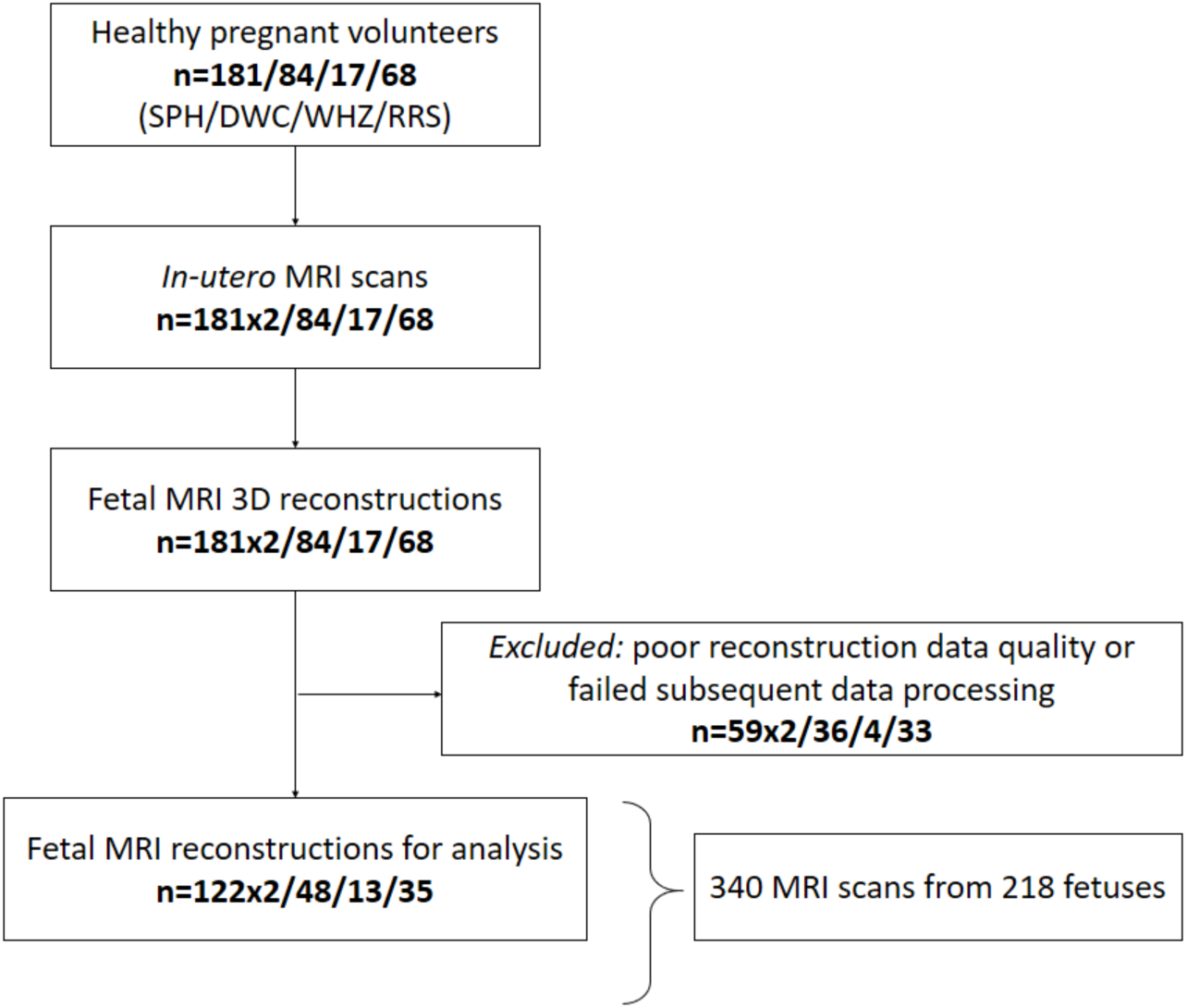
Flowchart shows inclusion and exclusion. SPH = Shandong Provincial Hospital, DWC = Dalian Municipal Women and Children’s Medical Center, WHZ = Women’s Hospital of Zhejiang University Medical School, RSS = Sir Run Run Shaw Hospital. Note that each fetus in SPH underwent two sets of MRI scans at 2 mm and 4 mm slice thickness.

**Figure S2.**
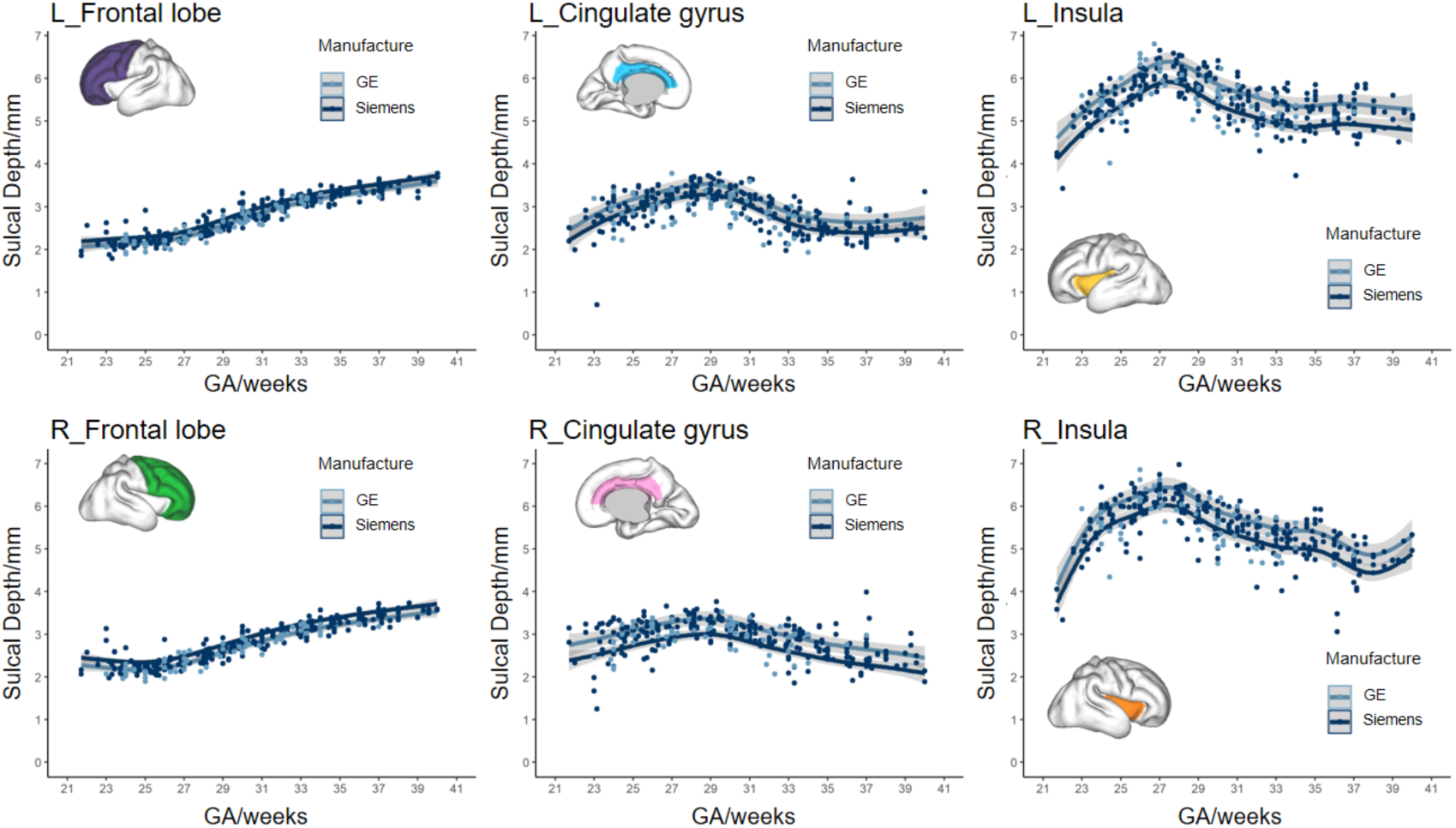
Effects of Manufacture (GE in light blue and Siemens in darker blue) on sulcal depth of left and right Frontal lobe (left column), Cingulate gyrus (middle column), and insula (right column).

## Reference

1. Hosny IA, Elghawabi HS (2010): Ultrafast MRI of the fetus: an increasingly important tool in prenatal diagnosis of congenital anomalies. Magn Reson Imaging. 28:1431–1439.

2. De Asis-Cruz J, Limperopoulos C (2022): Harnessing the Power of Advanced Fetal Neuroimaging to Understand In Utero Footprints for Later Neuropsychiatric Disorders. Biol Psychiatry.

3. Ebner M, Wang G, Li W, et al. (2020): An automated framework for localization, segmentation and super-resolution reconstruction of fetal brain MRI. Neuroimage. 206:116324.

4. Clouchoux C, Kudelski D, Gholipour A, et al. (2012): Quantitative in vivo MRI measurement of cortical development in the fetus. Brain Structure and Function. 217:127–139.

5. Habas PA, Scott JA, Roosta A, et al. (2011): Early Folding Patterns and Asymmetries of the Normal Human Brain Detected from in Utero MRI. Cereb Cortex. 22:13–25.

6. Rajagopalan V, Scott J, Habas PA, et al. (2011): Local Tissue Growth Patterns Underlying Normal Fetal Human Brain Gyrification Quantified In Utero. J Neurosci. 31:2878–2887.

7. Clouchoux C, Du Plessis AJ, Bouyssi-Kobar M, et al. (2013): Delayed Cortical Development in Fetuses with Complex Congenital Heart Disease. Cereb Cortex. 23:2932–2943.

8. Yun HJ, Lee HJ, Lee JY, et al. (2022): Quantification of sulcal emergence timing and its variability in early fetal life: hemispheric asymmetry and sex difference. Neuroimage.119629.

9. Machado-Rivas F, Gandhi J, Choi JJ, et al. (2021): Normal Growth, Sexual Dimorphism, and Lateral Asymmetries at Fetal Brain MRI. Radiology.

10. Button KS, Ioannidis JPA, Mokrysz C, et al. (2013): Power failure: why small sample size undermines the reliability of neuroscience. Nature Reviews Neuroscience. 14:365–376.

11. Van Essen DC, Ugurbil K, Auerbach E, et al. (2012): The Human Connectome Project: a data acquisition perspective. Neuroimage. 62:2222–2231.

12. Thompson PM, Jahanshad N, Ching CRK, et al. (2020): ENIGMA and global neuroscience: A decade of large-scale studies of the brain in health and disease across more than 40 countries. Translational Psychiatry. 10:100.

13. Sudlow C, Gallacher J, Allen N, et al. (2015): UK Biobank: An Open Access Resource for Identifying the Causes of a Wide Range of Complex Diseases of Middle and Old Age. PLoS Med. 12:e1001779.

14. Radua J, Vieta E, Shinohara R, et al. (2020): Increased power by harmonizing structural MRI site differences with the ComBat batch adjustment method in ENIGMA. Neuroimage. 218:116956.

15. Takao H, Hayashi N, Ohtomo K (2011): Effect of scanner in longitudinal studies of brain volume changes. J Magn Reson Imaging. 34:438–444.

16. Jovicich J, Czanner S, Han X, et al. (2009): MRI-derived measurements of human subcortical, ventricular and intracranial brain volumes: Reliability effects of scan sessions, acquisition sequences, data analyses, scanner upgrade, scanner vendors and field strengths. Neuroimage. 46:177–192.

17. Reig S, Sánchez-González J, Arango C, et al. (2009): Assessment of the increase in variability when combining volumetric data from different scanners. Hum Brain Mapp. 30:355–368.

18. Fortin J-P, Cullen N, Sheline YI, et al. (2018): Harmonization of cortical thickness measurements across scanners and sites. Neuroimage. 167:104–120.

19. Han X, Jovicich J, Salat D, et al. (2006): Reliability of MRI-derived measurements of human cerebral cortical thickness: The effects of field strength, scanner upgrade and manufacturer. Neuroimage. 32:180–194.

20. Auzias G, Takerkart S, Deruelle C (2016): On the Influence of Confounding Factors in Multisite Brain Morphometry Studies of Developmental Pathologies: Application to Autism Spectrum Disorder. IEEE Journal of Biomedical and Health Informatics. 20:810–817.

21. Dickerson BC, Fenstermacher E, Salat DH, et al. (2008): Detection of cortical thickness correlates of cognitive performance: Reliability across MRI scan sessions, scanners, and field strengths. Neuroimage. 39:10–18.

22. Liu M, Lepage C, Kim SY, et al. (2021): Robust Cortical Thickness Morphometry of Neonatal Brain and Systematic Evaluation Using Multi-Site MRI Datasets. Front Neurosci. 15.

23. Grigorescu I, Vanes L, Uus A, et al. (2021): Harmonized Segmentation of Neonatal Brain MRI. Front Neurosci. 15.

24. Kurokawa R, Kamiya K, Koike S, et al. (2021): Cross-scanner reproducibility and harmonization of a diffusion MRI structural brain network: A traveling subject study of multi-b acquisition. Neuroimage. 245:118675.

25. Wang Y-W, Chen X, Yan C-G (2023): Comprehensive evaluation of harmonization on functional brain imaging for multisite data-fusion. Neuroimage.120089.

26. Noble S, Scheinost D, Finn ES, et al. (2017): Multisite reliability of MR-based functional connectivity. Neuroimage. 146:959–970.

27. Jovicich J, Marizzoni M, Bosch B, et al. (2014): Multisite longitudinal reliability of tract-based spatial statistics in diffusion tensor imaging of healthy elderly subjects. Neuroimage. 101:390–403.

28. Onicas AI, Ware AL, Harris AD, et al. (2022): Multisite Harmonization of Structural DTI Networks in Children: An A-CAP Study. Front Neurosci. 13.

29. Chu DY, Adluru N, Nair VA, et al. (2023): Application of data harmonization and tract-based spatial statistics reveals white matter structural abnormalities in pediatric patients with focal cortical dysplasia. Epilepsy Behav. 142:109190.

30. Zhong J, Wang Y, Li J, et al. (2020): Inter-site harmonization based on dual generative adversarial networks for diffusion tensor imaging: application to neonatal white matter development. Biomed Eng Online. 19:4.

31. Johnson WE, Li C, Rabinovic A (2007): Adjusting batch effects in microarray expression data using empirical Bayes methods. Biostatistics. 8:118–127.

32. Fortin JP, Parker D, Tunç B, et al. (2017): Harmonization of multi-site diffusion tensor imaging data. Neuroimage. 161:149–170.

33. Yu M, Linn KA, Cook PA, et al. (2018): Statistical harmonization corrects site effects in functional connectivity measurements from multi-site fMRI data. Hum Brain Mapp. 39:4213–4227.

34. Pomponio R, Erus G, Habes M, et al. (2020): Harmonization of large MRI datasets for the analysis of brain imaging patterns throughout the lifespan. Neuroimage. 208:116450.

35. Saleem SN (2014): Fetal MRI: An approach to practice: A review. Journal of Advanced Research. 5:507–523.

36. Avants BB, Epstein CL, Grossman M, et al. (2008): Symmetric diffeomorphic image registration with cross-correlation: evaluating automated labeling of elderly and neurodegenerative brain. Med Image Anal. 12:26–41.

37. Gholipour A, Rollins CK, Velasco-Annis C, et al. (2017): A normative spatiotemporal MRI atlas of the fetal brain for automatic segmentation and analysis of early brain growth. Sci Rep. 7:476–476.

38. Isensee F, Jaeger PF, Kohl SAA, et al. (2021): nnU-Net: a self-configuring method for deep learning-based biomedical image segmentation. Nat Methods. 18:203–211.

39. Zhou Z, Rahman Siddiquee MM, Tajbakhsh N, et al. (2018): UNet++: A Nested U-Net Architecture for Medical Image Segmentation. In: Stoyanov D, Taylor Z, Carneiro G, Syeda-Mahmood T, Martel A, Maier-Hein L, et al., editors. Deep Learning in Medical Image Analysis and Multimodal Learning for Clinical Decision Support. Cham: Springer International Publishing, pp 3–11.

40. Çiçek Ö, Abdulkadir A, Lienkamp SS, et al. (2016): 3D U-Net: Learning Dense Volumetric Segmentation from Sparse Annotation. In: Ourselin S, Joskowicz L, Sabuncu MR, Unal G, Wells W, editors. Medical Image Computing and Computer-Assisted Intervention – MICCAI 2016. Cham: Springer International Publishing, pp 424–432.

41. Makropoulos A, Gousias IS, Ledig C, et al. (2014): Automatic Whole Brain MRI Segmentation of the Developing Neonatal Brain. IEEE Trans Med Imaging. 33:1818–1831.

42. Makropoulos A, Robinson EC, Schuh A, et al. (2018): The developing human connectome project: A minimal processing pipeline for neonatal cortical surface reconstruction. Neuroimage. 173:88–112.

43. Xinyi X, Cong S, Jiwei S, et al. (2022): Spatiotemporal Atlas of the Fetal Brain Depicts Cortical Developmental Gradient. The Journal of Neuroscience. 42:9435.

44. Wood SN (2006): Generalized additive models: an introduction with R. chapman and hall/CRC.

45. Benjamini Y, Hochberg Y (1995): Controlling the False Discovery Rate: A Practical and Powerful Approach to Multiple Testing. 57:289–300.

46. Yin X, Goudriaan J, Lantinga EA, et al. (2003): A Flexible Sigmoid Function of Determinate Growth. Ann Bot. 91:361–371.

47. Pfefferbaum A, Rohlfing T, Rosenbloom MJ, et al. (2012): Combining atlas-based parcellation of regional brain data acquired across scanners at 1.5T and 3.0T field strengths. Neuroimage. 60:940–951.

48. Lee H, Nakamura K, Narayanan S, et al. (2019): Estimating and accounting for the effect of MRI scanner changes on longitudinal whole-brain volume change measurements. Neuroimage. 184:555–565.

49. Biberacher V, Schmidt P, Keshavan A, et al. (2016): Intra- and interscanner variability of magnetic resonance imaging based volumetry in multiple sclerosis. Neuroimage. 142:188–197.

50. Gebre RK, Senjem ML, Raghavan S, et al. (2023): Cross–scanner harmonization methods for structural MRI may need further work: A comparison study. Neuroimage. 269:119912.

51. Fennema-Notestine C, Gamst AC, Quinn BT, et al. (2007): Feasibility of Multi-site Clinical Structural Neuroimaging Studies of Aging Using Legacy Data. Neuroinformatics. 5:235–245.

52. Hu F, Chen AA, Horng H, et al. (2023): Image harmonization: A review of statistical and deep learning methods for removing batch effects and evaluation metrics for effective harmonization. Neuroimage. 274:120125.

53. Sun D, Rakesh G, Haswell CC, et al. (2022): A Comparison of Methods to Harmonize Cortical Thickness Measurements Across Scanners and Sites. Neuroimage.119509.

